## Supporting information for "On the Quina side: A Neanderthal bone industry at Chez-Pinaud site, France"

**S1 Fig. Archaeological deposits of the Chez-Pinaud site (Jonzac, France).** (a) location of the excavation area in 2019-2021. (b) left stratigraphic cut (after Airvaux and Soressi 2005). (c) view of the "bone-bed", 2021 excavation (photos: W. Rendu).

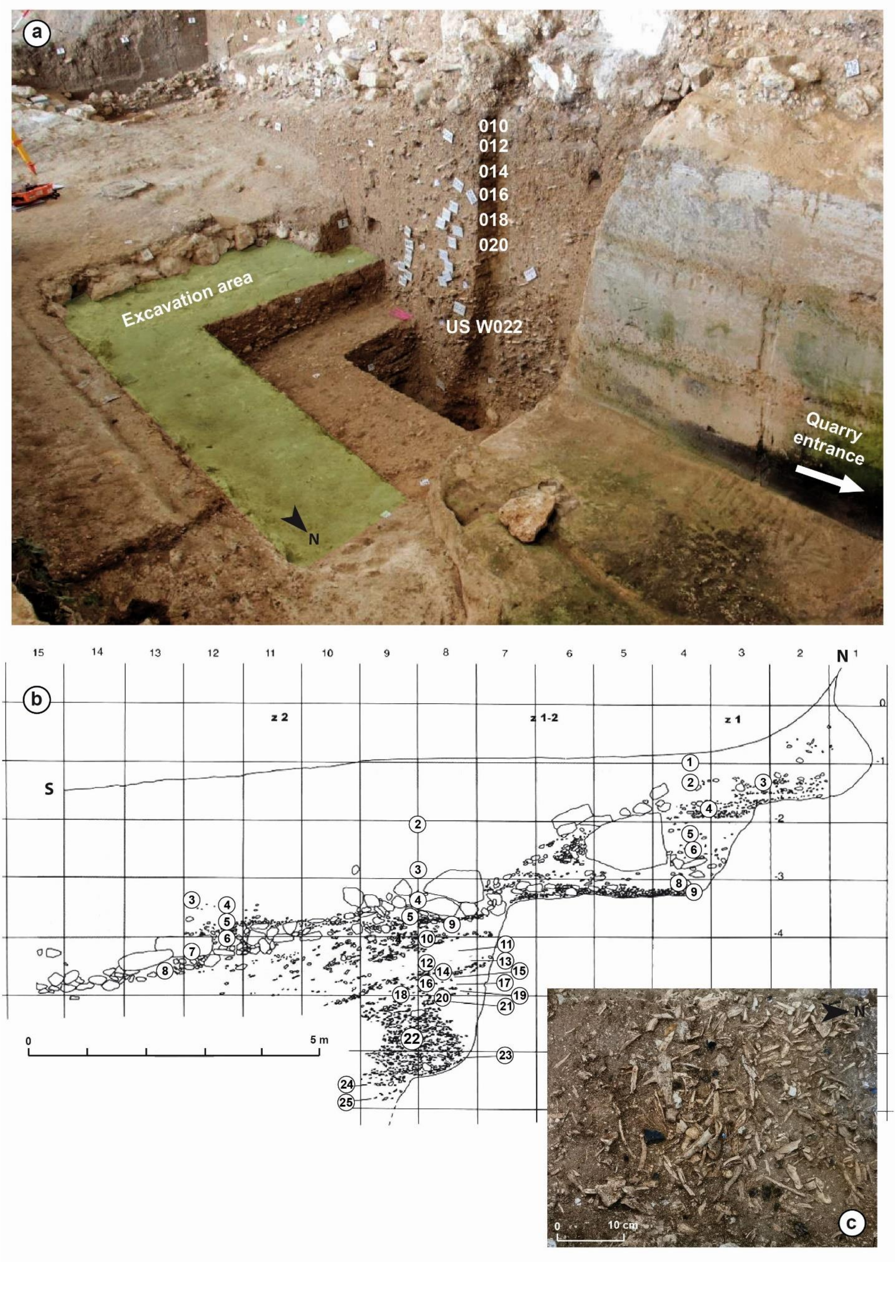

**S2 Fig. Experiments with replicas of Mousterian type bone tools. (**a) fracturation by direct percussion on anvil; (b) fractured fresh long bones of Bovinae. (c) regular flakes from bone fracturing (d) impact of direct percussion. (e) retouched bone blank. (f) flake from retouch. (g) Mousterian side scrapers shaping. (h) meat cutting (i) hair removing from skin. (j) wooden peeling. (k) plant harvesting. (l) soil digging (photos: H. Plisson).

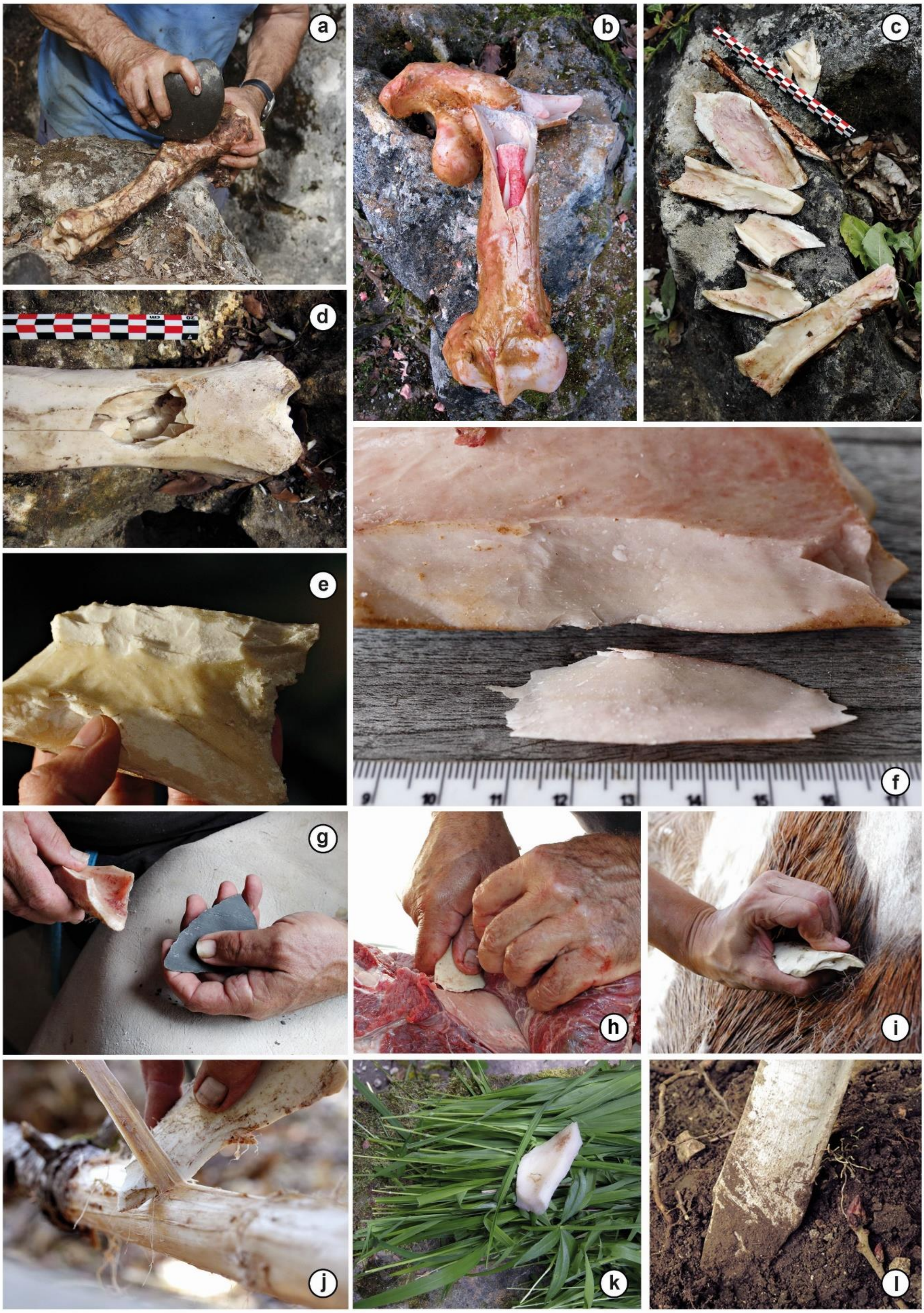

**S3 Fig. Examples of ribs with smoothed end discovered in other Middle Paleolithic contexts.** (a), (c–d) Abri Peyrony. (b) Pech de l'Azé (after Soressi et al. 2013). (e) Zaskalnaya VI (after Stepanchuk et al. 2017). (f) Chagyrskaya Cave (after Baumann et al. 2020). (g) Abri des Canalettes (after Patou-Mathis 1993). (h) Axlor (after Mozota Holgueras 2012). (i–k) La Quina (after Henri-Martin 1907–1910). (l–m) Grotte du Noisetier (after Oulad El kaïd 2016).

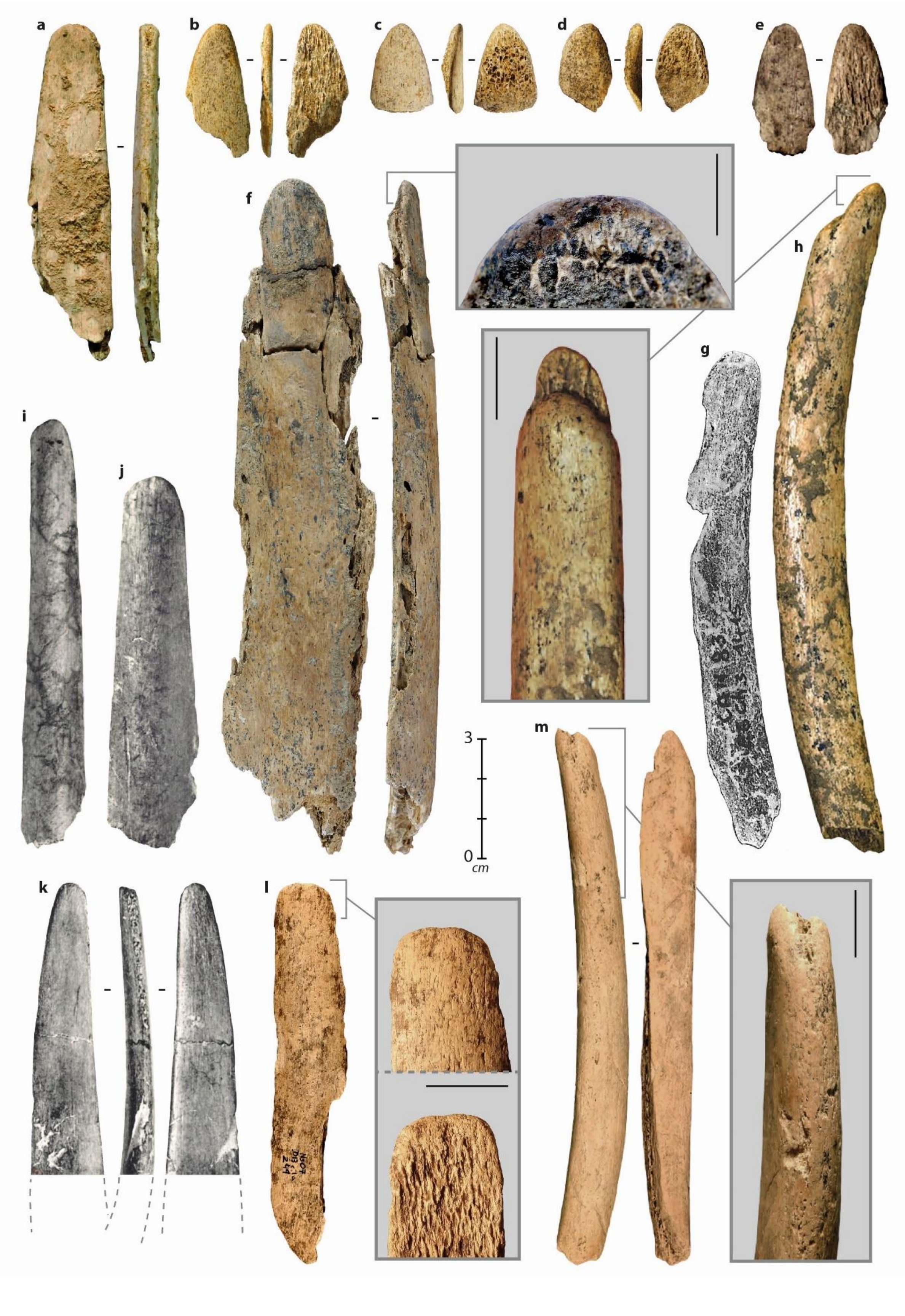

**S4 Fig. Examples of retouched bone artifacts discovered in pre-AMH contexts.** (a) Castel di Guido (after Villa et al. 2021). (b) Nova de Columbeira (after Zilhão et al. 2011). (c) Vauffrey (after Vincent 1993). (d) Poggeti Vecchi (after Aranguren et al. 2019). (e) Bois-Roche (after Vincent 1993). (f) Combe-Grenal (after Tartar and Costamagno 2016). (g–h) Chagyrskaya (photo: M. Baumann). (i) Gran Dolina (after Rossel et al. 2011). (j) Abric Romaní (after Carbonel et al. 1994).

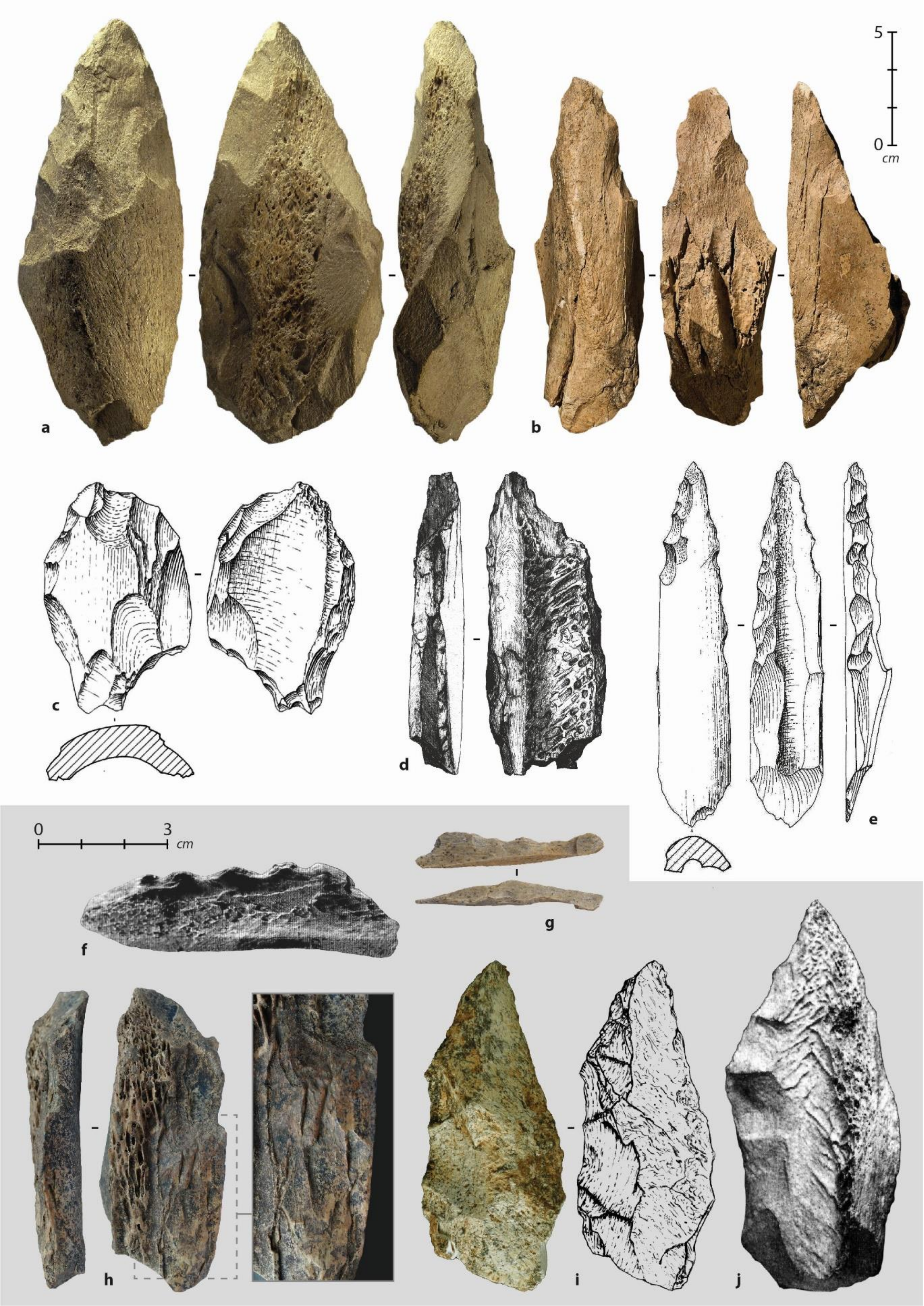

### S1 Table. Microtomographic recording parameters of the experimental and archaeological tools from Chez-Pinaud site.

| | Identification number | Raw material | Tool type | Scanned area | Resolution (μm) | Slice nb. | Correction value | kV | μA |
|---|---|---|---|---|---|---|---|---|---|
| Archaeological tools | CPN19-5 | Diaphysis large size ungulate | Retouched | Complete | 26 | 1625 | 10.40 | 120 | 120 |
| | CPN19-529 | Diaphysis medium size ungulate | Retouched | Complete | 26 | 1625 | 10.40 | 120 | 120 |
| | CPN19-534 | Humerus large size ungulate | Beveled | Complete | 35 | 2125 | 8.32 | 120 | 120 |
| | CPN19-888 | Diaphysis large size ungulate | Beveled Retouched | Complete | 35 | 1500 | 9.13 | 120 | 120 |
| | CPN19-1014 | Horse humerus | Beveled Retouched Retoucher | Complete | 67 | 1200 | 2.46 | 120 | 250 |
| | | | | ROI* 1 | 21 | 2550 | 3.52 | 120 | 250 |
| | CPN19-2020 | Horse tibia | Retoucher | Complete | 62 | 1200 | 2.48 | 120 | 250 |
| | | | | ROI 1 | 22 | 2250 | 4.49 | 120 | 250 |
| | CPN19-2132 | Diaphysis large size ungulate | Beveled ? Retouched Retoucher | Complete | 43 | 1375 | 7.12 | 120 | 120 |
| | CPN20-3581 | Rib large size ungulate | Smoothed | Complete | 54 | 1275 | 8.31 | 120 | 100 |
| | | | | ROI 1 | 14 | 1625 | 27.10 | 120 | 100 |
| | | | | ROI 2 | 5 | 2550 | 74.83 | 120 | 120 |
| | CPN20-3609 | Bison metatarsal | Beveled Smoothed | Complete | 43 | 1375 | 8.41 | 120 | 120 |
| Experimental tools | Exp-10 | Cow tibia | Beveled | Complete | 89 | 1200 | 2.82 | 120 | 250 |
| | | | | ROI 1 | 13 | 2250 | 8.24 | 120 | 250 |
| | | | | Zone 2 | 13 | 2250 | 1.69 | 120 | 250 |
| | Exp-13 | Red deer femur | Beveled Retouched | Complete | 89 | 1200 | 2.82 | 120 | 250 |
| | | | | ROI 1 | 24 | 2250 | 5.22 | 120 | 250 |
| | | | | ROI 2 | 24 | 2250 | 2.12 | 120 | 250 |
| | Exp-46 | Cow tibia | Retoucher Retouched | Compete | 98 | 1500 | 2.78 | 120 | 250 |
| | | | | ROI 1 | 21 | 2250 | 6.18 | 120 | 250 |
| | Exp-57 | Cow tibia | Retoucher | Complete | 98 | 1200 | 3.18 | 120 | 250 |
| | | | | ROI 1 | 21 | 2550 | 3.22 | 120 | 250 |
| | Exp-58 | Cow tibia | Retoucher | Complete | 98 | 1200 | 3.18 | 120 | 250 |
| | | | | ROI 1 | 33 | 2250 | 4.02 | 120 | 250 |

*ROI = Region of interest

#### S2 Table. Experimental samples analyzed in µCT.

| | **Exp-10** | **Exp-13** | **Exp-46** | **Exp-57** | **Exp-58** |
|---|---|---|---|---|---|
| Raw material | Cow Tibia | Red deer Femur | Cow Tibia | Cow Tibia | Cow Tibia |
| Time after death (months) | 4 | 10 | 0 | 1 | 15 |
| 1st fracturing step | Direct percussion/ on anvil | Direct percussion/ on anvil | Direct percussion/ on anvil | Direct percussion/ on anvil | Direct percussion/ on anvil |
| Shaping | None | Retouched | Retouched | None | None |
| Shaping technique | None | Direct percussion/ Soft hammer | Direct percussion/ Soft hammer | None | None |
| Tool type | Beveled | Beveled | Retoucher | Retoucher | Retoucher |
| Length (cm) | 15.9 | 13.21 | 16.84 | 11.47 | 18.1 |
| Width (cm) | 3.27 | 4.41 | 5.97 | 3.61 | 4.96 |
| Thickness (cm) | 1.24 | 0.71 | 1.29 | 1.54 | 1.23 |
| Worked material | Fresh *Corylus avellana* | Fresh *Corylus avellana* | “Bergerac” flint | “Bergerac” flint | “Bergerac” flint |
| Activity | Handle manufacture | Handle manufacture | Scraper shaping | Scraper shaping | Scraper shaping |
| Task | Branch hollowing | Branch hollowing | Edge retouching | Edge retouching | Edge retouching |
| Time (mn.) | 20 | 20 | < 5 | < 5 | < 5 |

**S3 Table. Reports (non-exhaustive) of smoothed-end ribs from Neanderthal contexts in Eurasia.**

| Loc | Site | Nb. | Industry | Dates | Ref. |
|---|---|---|---|---|---|
| France | Abri Peyrony | 3 | MTA | MIS 3 (43 – 37 ky BP) | Soressi et al. 2013 |
| | Noisetier | 2 | Discoid | MIS 3 (42 ky BP) | Oulad El Kaïd 2016 |
| | La Quina | 3 | MAT/Denticulate | MIS 3 (48 - 40 ky BP) | Henri-Martin 1907-1910<br>Debénath et al. 1998 |
| | Lartet | 4 | Levallois | MIS 3 (48 - 35 ky BP) | Debénath and Duport 1971 |
| | Pech de l'Azé | 1 | MTA | MIS 3 (51 ky BP) | Soressi et al. 2013 |
| | Pradayrol | 1 | Discoid<br>Levallois | MIS 3 | Villeneuve et al. 2019 |
| | Canalettes | 1 | Levallois | MIS 5-4 (73 ky BP) | Patou-Mathis 1993<br>Valladas et al. 1987 |
| | Vaufrey | 1 | Mousterian | MIS 7-6 (270-142 ky BP) | Vincent 1993 |
| Spain | Cueva Mórin | 1 | MTA (Vasconian) | MIS 3 (>43 ky BP) | Freeman 1971<br>Maíllo-Fernández et al. 2014 |
| | Axlor | 1 | Quina | MIS 3 (>47 ky BP) | Mozota Holgueras 2012<br>Gómez-Olivencia et al. 2018 |
| Germany | Salzgitter-Lebenstedt | 8 | Levallois | MIS 3 (55 – 48 ky BP) | Gaudzinski 1999<br>Pastoors 2009 |
| Crimea | Zaskalnaya VI | 1 | Levallois (Ak-Kaya) | MIS 3 (39 – 30 ky BP) | Stepanchuk et al. 2017 |
| Siberia | Chagyrskaya | 2 | Micoquian (Sibiryachika) | MIS 4-3 (60 – 50 ky BP) | Baumann et al. 2020 |

### S4 Table. Reports (non-exhaustive) of knapped bone tools from pre-AMH contexts in Eurasia.

| Loc | Site | Nb. | Industry | Dates | Ref. |
|---|---|---|---|---|---|
| France | Combe-Grenal | 1<br>1 | Mousterian Levallois<br>Denticulate | MIS 3 (39-38 ky BP) | Bordes 1961<br>Vincent 1993<br>Tartar and Cotamagno 2016 |
| | Jonzac | 7 | Quina | MIS 4 (72 ky BP) | Rendu et al. 2020<br>Richter et al. 2013 |
| | Vaufrey | 1 | Mousterian Levallois | MIS 4 (74 ky BP) | Vincent 1993<br>Tartar and Costamagno 2016 |
| | Pié-Lombard | 4 | Levallois | MIS 5 (70 ky BP) | Texier 1974<br>Texier et al. 2011 |
| | Rigabe | 6 | Levallois | MIS 3-5 | Defleur 1988<br>Brugal et al. 2020 |
| | La Ferrassie | 1 | Mousterian | NR | Bordes, 1961 |
| | Bois-Roche | +/-15 | Mousterian | NR | Vincent 1993 |
| | Baume de Gigny | 1 | Mousterian | NR | Vuillemey 1989 |
| Germany | Rhede | 1 | Micoquian (Keilmesser) | MIS 5 (70 ky BP) | Tromnau 1983<br>Baales and Stapel 2015 |
| | Sirgenstein | 1 | Mousterian | NR | Hahn 1976<br>Ono 2006 |
| Belgium | Trou Magrite | 1 | Mousterian | MIS 3 | Personal inventory<br>Jimenez et al. 2016 |
| Italy | Fumane | 1 | Levallois | MIS 3 (42 ky BP) | Romandini et al. 2014<br>Peresani et al. 2013 |
| | Poggeti Vecchi | 10 | Mousterian | MIS 6-7 (171 ky BP) | Aranguren et al. 2019 |
| | Casal de'Pazzi | 1 | Protopontinian | MIS 7 (270-250 ky BP) | Anzidei and Gioia 1992<br>Marra et al. 2018<br>Villa et al. 2021 |
| | La Polledrara | +/-3 | Acheulean (without bifaces) | MIS 9 (324 ky BP) | Santucci et al. 2016<br>Villa et al. 2021 |
| | Lademagne | +/-2 | Acheulean | MIS 10-11 (405-389 ky BP) | Pereira et al. 2018<br>Villa et al. 2021 |
| | Castel di Guido | 81 | Acheulean | MIS 11 (395 ky BP) | Boschian and Saccà 2015<br>Villa et al. 2021 |

| | | | | | |
|---|---|---|---|---|---|
| | Fontana Ranuccio | 5 | Acheulean | MIS 11 (407 ky BP) | Biddittu and Serge 1982<br>Pereira et al. 2018<br>Villa et al. 2021 |
| | Malagrotta | +/-6 | Acheulean (without bifaces) | MIS 11 (451-378 ky BP) | Marra and Gatta 2019<br>Villa et al. 2021 |
| | Pontecorvos | 1 | Acheulean | NR | Biddittu and Serge 1982<br>Villa et al. 2021 |
| Spain | Axlor | 6 | Quina | MIS 3 (>47 ky BP) | Mozota Holgueras 2012<br>Baldéon 1999 |
| | Peña Miel | +/- 22 | Quina | MIS 3 (+/-50 ky BP) | Barandiarán 1987<br>Montes et al. 2001 |
| | Abric Romaní | 1 | Denticulate | MIS 4-3 (61-39 ky BP) | Tartar and Costamagno 2016<br>Carbonel et al. 1994 |
| | Gran Dolina | 2 | Mousterian | MIS 9 (372-244 ky BP) | Rossel et al. 2011 |
| | Bolomor | 1 | Denticulate | MIS 9 (350 ky BP) | Rossel et al. 2015 |
| Portugal | Nova de Columbeira | 1 | Mousterian | (87 ky BP) | Zilhão et al. 2011 |
| Czech Republic | Kůlna | 42 | Micoquian | MIS 3 (125-45 ky BP) | Vincent 1993 |
| | | 27 | Taubachian | | |
| | | 4 | Levallois | | |
| Russia | Denisova | 4 | Levallois (C 11.4) | MIS 5 (105 ky BP) | Jacobs et al. 2019<br>Kozlikin et al. 2020 |
| | Chagyrskaya | 49 | Micoquian (Sibiryachika) | MIS 4-3 (60 – 50 ky BP) | Baumann et al. 2020 |
| Japan | Tategahana | > 3 | NR | MIS 3 | Ono 2006<br>Nojiri group 1985 |
| | Jinniushan Loc A | NR | NR | (230-200 ky BP) | Ono 2006 |

### S1 Text. Bone retouchers

The bone retoucher is a common Middle Paleolithic bone tool (Armand and Delagnes 1998; Mallye et al. 2012; Jequier et al. 2012; Mozota Holgueras 2012; Blasco et al. 2013; Abrams et al. 2014; Deaujard et al. 2014; Rosell et al. 2015; Rougier et al. 2016; Doyon et al. 2018; Costamagno et al. 2018; Mateo-Lomba et al. 2019), which already exist in previous periods (Roberts and Parfitt 1999; Langlois 2004; Smith 2013; Julien et al 2015; Kolfschoten et al. 2015; Moigne et al. 2016) and last throughout the Upper Paleolithic (Patou-Mathis 2002; Castel et al 2003; Castel and Madelaine 2003; Rigaud 2007; Tartar 2009, 2012). Identified from its use-wear traces, it has benefited from functional analyses since the first specimens were discovered (Henri-Martin 1906). Most scholars agree that it is a light hammer for shaping lithic edges (Bonch-Osmolovskyi 1940; Semenov 1964; Feustel 1973; Rigaud 1977, 2007; Schelinskii 1983; Vincent 1988; Bourginon 2001; Schwab 2009; Mozota 2012; Mallye et al. 2012). But bone retouchers are seen as an undifferentiated unit, despite their morphometrical diversity, because of the scores apparent uniformity that does not allow to see significant differences between sites or chronocultures, apart from of the longitudinal orientation of the scores, specific to Upper Paleolithic material (Schwab 2002).

#### S2 Text. Beveled tools

Beveled tools, mainly made from antler, are frequently identified in Upper Paleolithic contexts (Deffarges et al. 1974; Provenzano 1984). In later periods, Mesolithic and Neolithic, bone specimens are more common (Camps-Fabrer et al. 1998; Maigrot 2003). In techno-traceological studies, chisels (for cutting) and wedges (for splitting) are often grouped together (Provenzano 1998) because they share the same characteristics: a cutting edge at one end marked by crushing sometimes associated with a blunt area, chips or small removals, and striations oriented along the main axis of the tool. The force imparted to set them in motion can be transmitted directly by the arm, or indirectly with a hammer. In the latter case, the tools are characterized by a striking surface located on the end opposite to the bevel and materialized by a crushing area surrounded by macro to micro removals. However, the absence of a striking surface does not necessarily indicate that a hammer has not been used. This occurs when the tools are hafted, such as the bone bevels from the Swiss Final Neolithic (Voruz 1984). If the handle is not preserved, it can be revealed by: (1) the proximal end shaping (Sidéra 1989), (2) a glossing/blunting of the proximal tool edges (Maigrot 1997), a clear limit of the distal use-wear traces, or a clear change of the bone material coloration (Maigrot 2003). Repeated hammering of the proximal end can also lead to fatigue fractures. These use-fractures are reportedly quite common on bone beveled tools (Tartar 2012; Maigrot et al. 2013).

#### S3 Text. Retouched tools

The identification of an intentional retouch on bone is often put forward when all other causes that could lead to removals have been ruled out (Vincent 1993). The archaeological context allows the diagnosis to be based with greater confidence (Inizan et al. 1995). At Chez-Pinaud, the absence of carnivore consumption traces (chewing, grooves, pits, punctures, and digestion blunts; Stucliffe 1973; Haynes 1983; Campas and Beauval 2008; Fourvel 2012), dispels the first source of possible confusion regarding the notches origin. The likelihood that the bone blanks were transformed by humans for technical purposes is here increased by their tool status: most were used as retouchers. Among the technical causes, bone fracturing by percussion during butchery processes also lead to the formation of removals (medullary or cortical side). In this case, the latter are related to the adjacent fracture, and sometimes to traces of the anvil's counter-strike on the opposite face, all resulting from a single event, generally a violent blow given perpendicular to the bone (Capaldo and Blumenschine 1994; Pickering and Egeland 2006).

Although there are few references about the intentional retouch of bone, positive clues of identification can nevertheless be used. Lithic knapping provides keys to understanding that can partially be transferred to bone insofar as the latter behaves like a conchoidal fracture material. However, bone is also a fibrous material whose mechanical properties vary according to the stress direction. Therefore, it responds to knapping in a more complex way than flint, due to its anisotropic structure. Nevertheless, common regularities and characteristics can be observed, implying the same reasoning in their analysis. The organization of the removals is the main criterion to consider for identifying an intentional knapping (Lyman 1984; Vincent 1993; Inizan et al. 1995).

#### S4 Text. Smooth-ended tools

Blunts are among the most common bone alterations induced by a variety of post-depositional processes (Sutcliffe 1973; Shipman and Rose 1988; Olsen 1989; Villa and d'Errico 2001; Fernández-Jalvo and Andrews 2013). The main criteria for differentiating a technical blunt from a taphonomic one is its extension, location, orientation, its transition with adjacent surfaces, and the combination of these different features. Natural erosion or dissolution tends to cover most, if not all, of the bone surface, whereas a blunt from use is restricted to the surface or edge of the active area. Hafting or gripping blunts may be more extensive, but remain related to a particular tool area. Blunt from friction into the sediments is mainly superficial and does not affect the concavities. The tool effectiveness depends on the control of its working angle. Maintaining this angle induces a regularity of the use-wear characteristics. The profile of the active edge rounding (seen in cross section) thus depends on the way the tool is used but also on the worked material physical properties, depending on whether the latter is organic or mineral, dry, fresh or wet, compact or granular, fibrous or not, etc. The maximum development of the bluntness is fixed by the minimum sharpness necessary to obtain the expected results on the worked material.
